# ASVNet: Inferring microbes from *16S rRNA* amplicon sequencing data

**DOI:** 10.1101/2025.11.21.689717

**Authors:** Juan J. Sánchez-Gil, Ronnie de Jonge

## Abstract

Amplicon sequencing of the variable regions of the *16S rRNA* gene is one of the most common techniques in microbiome research. A crucial step in analysing this data is to cluster raw sequences into biologically interpretable groups that approximate the actual organisms in a community, which reduces data complexity, facilitates diversity analyses, and enables clearer ecological and functional interpretations. Most approaches rely on fixed sequence similarity thresholds, typically operational taxonomic units (OTUs) at 97% or amplicon sequence variants (ASVs) at 99-100%. However, finding the correct correspondence between amplicons and the original organisms in a complex community remains challenging, primarily due to the presence of varying copy numbers within genomes and the high similarity of amplicon sequences among species. While the stringent threshold in ASVs prevents merging sequences from different species, it often splits sequences from the same genome into multiple ASVs, inflating species counts. Conversely, the 97% similarity threshold in OTUs tends to frequently collapse multiple distinct organisms into a single unit regardless of their functional or ecological (dis)similarity, and thereby underestimating true diversity. To address these limitations, we introduce ASVNet, an algorithmic framework that leverages amplicon sequence similarity and co-abundance networks to better reflect the underlying organisms in a community. ASVNet builds on the principle that amplicon counts from the same organism co-vary across samples and it thus attempts to cluster these co-occurring, similar sequences as empirical OTUs (eOTUs). Comparative analyses in synthetic and natural root-associated microbial communities with genome-sequenced isolates show that ASVNet-derived eOTUs more accurately represent the original microbes and possess greater biological relevance than OTUs defined at any fixed threshold. By providing a biologically grounded framework to amplicon sequence data interpretation, ASVNet extends our ability to extract meaningful insights from complex microbiome datasets.

## Main

### Challenges in amplicon sequencing for microbial community analysis

In prokaryotes, the genes responsible for the production of ribosomal RNA (rRNA) are commonly contained within the *rrn* operon, although it is frequent to find individual genes located elsewhere in the genome (Brewer et al., 2020). Due to the high amounts of rRNA required for cell survival and reproduction, the *rrn* operon is found in multiple copies in 92% of known bacteria, where it generally ranges from 1 to 37 copies per genome (Pan et al., 2023; Větrovský and Baldrian, 2013). The *rrn* copy number varies according to taxonomy and determines key (eco)physiological traits, such as maximum growth rate, overall metabolic capacities, and ability to respond to fluctuating and complex environments (Klappenbach et al., 2000; Lin et al., 2023; Roller et al., 2016). Nevertheless, this essential role of *rrn* genes in bacterial metabolism imposes strong evolutionary restrictions, which translates in high conservation of both its sequence and structure (Pei et al., 2010), a property that has enabled their extensive use as a phylogenetic marker for decades (Goossens et al., 2023; Johnson et al., 2019; Winker and Woese, 1991).

Particularly, the *rrnB* gene, also commonly referred to as *16S rRNA* (hereafter, *16S* gene), encodes the 1,5 kbp long *16S rRNA* that conforms the nucleotidic portion of the small ribosomal subunit. Despite having a highly conserved sequence, the *16S* gene harbours nine hypervariable regions (V1-V9) that exhibit taxon-specific variability throughout prokaryotic genomes. Consequently, amplicon sequencing of these hypervariable regions is the most common technique for microbial identification in microbiome research, although sequencing of the entire *16S* gene is becoming increasingly more popular as it enables enhanced taxonomic resolution.

In the archetypal microbiome study, the desired region of the *16S* gene is PCR-amplified with primers located at the conserved flanking regions. Subsequently, the amplicons are sequenced and the read counts for every sequence are recorded and processed to obtain a representation of the microbial community. However, the amplification and sequencing steps are prone to errors and the raw data accrues many spurious reads that emerge from technical imprecisions. Consequently, algorithmic efforts in microbiome data processing focus on removal and/or correction of these erroneous sequences, a procedure known as denoising (Callahan et al., 2016).

The complex nature of the *16S* gene introduces significant biological and technical challenges when analysing amplicon microbiome data. As mentioned above, bacterial genomes often carry multiple copies of the *16S* gene, causing an overrepresentation of organisms with higher copy number (Větrovský and Baldrian, 2013). Moreover, about 60% of prokaryotic genomes harbour multiple copies of the gene, which can differ in their sequence (intragenomic heterogeneity) (Bodilis et al., 2012; Coenye and Vandamme, 2003; Pan et al., 2023; Pei et al., 2010). Furthermore, identical or highly similar copies are commonly shared across different species or genera (intergenomic conservation), resulting in copies being more similar across genomes than within a single genome (Pan et al., 2023). Consequently, some amplicon sequences represent multiple organisms simultaneously, while others remain organism-specific, leading to skewed microbial abundance estimates.

Additionally, the average *16S* copy number and the degree of intra- and intergenomic sequence similarity vary across taxa, further distorting the final community representation (**Figure 1A**; Schloss, 2021; Větrovský and Baldrian, 2013).

**Figure 1.**
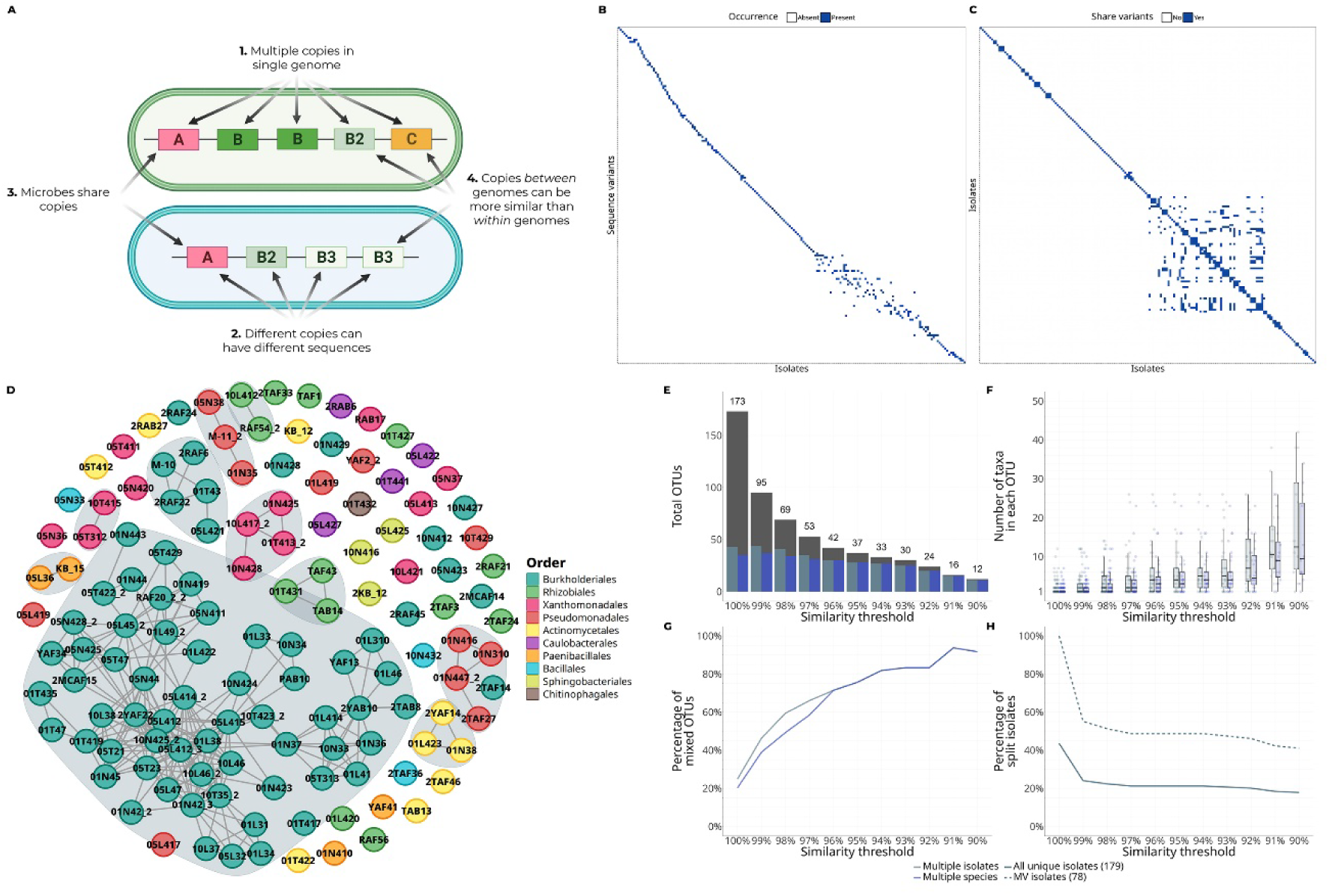
Single similarity thresholds alone cannot correctly cluster microbes. (A) Sources of complexity in *16S* gene amplicon sequencing datasets. All these phenomena increase the complexity in the analysis of this type of data, making it impossible to classify sequences in a biologically meaningful way based only on a fixed threshold. (B) Incidence matrix of the 134 isolates included in the SynCom and all *16S* gene sequence variants in their genomes. Cells in blue denote presence of a variant in an isolate genome. Note how the relationship variant-isolate is relatively simple in many isolates, but very complex in a group of them located at the bottom-right of the matrix. (C) Adjacency matrix of the isolates, depicting pair-wise co-occurrence of variants in their genomes. Blue cells denote events in which the same sequence variant occurs in both isolates in the comparison. The intricate set of interactions, seen towards the bottom-right of the matrix, indicate complex relationships between these isolates in terms of the variants that they share. (D) Network visualisation of the adjacency matrix in C, showing the isolates connected when they share V3-V4 sequences. The shaded areas enclose groups of isolates that share sequences between them but not with other isolates outside. (E) Total number of *k*OTUs found by clustering sequences from SynCom isolates at decreasing similarity thresholds. The blue-grey and blue bars show the number of those *k*OTUs that are chimeric at the isolate and species level, respectively. (F) Number of chimeric *k*OTUs containing either different isolates (blue-grey) and species (blue). (G) Percentage of chimeric *k*OTUs at the same similarity thresholds, depicting OTUs merging multiple isolates (blue-grey) and species (blue). (H) Percentage of isolates that are split into different *k*OTUs at different similarity thresholds, shown as a fraction of all isolates (full line) and of MV isolates (dashed line).

To date, numerous approaches have been developed to process and analyse *16S* amplicon data (Prodan et al., 2020). The two predominant strategies differ primarily in the similarity thresholds used for clustering. Amplicon sequence variants (ASVs) aim for high sequence resolution (99-100% similarity) and try to group identical sequences after denoising. While this prevents merging sequences from distinct species, ASVs often split multiple variants of the *16S* gene from the same genome into separate clusters (Schloss, 2021), artificially elevating diversity estimates. Consequently, ASV-based analyses tend to overestimate taxa with many distinct copies and underestimate those with fewer or shared copies (Pan et al., 2023). Alternatively, operational taxonomic units (OTUs) group sequences at a fixed similarity threshold (typically at 97% similarity), designed to encapsulate species-level variation (Konstantinidis and Tiedje, 2005). In contrast to ASVs, OTUs underestimate diversity by merging distinct organisms and species into chimeric clusters, regardless of their functional similarity or evolutionary relationships (Nguyen et al., 2016; Pan et al., 2023; Schloss, 2021). However, at the same time a threshold as low as 94.75% is required to merge all seven *16S* copies in *Escherichia coli* genomes into a single OTU (Schloss, 2021), highlighting the inherent limitations of fixed sequence similarity thresholds for cluster definition.

In summary, no single fixed similarity threshold fully captures the complex relationship between individual *16S* copies and their source organisms. Consequently, fixed thresholds negatively impact statistical analyses and biological interpretations by artificially inflating or underestimating diversity, splitting variants from single organisms, or merging distinct organisms into inaccurate clusters. We reasoned that these problems would be alleviated by correctly clustering amplicon sequences into the original organisms from which they arise. With this goal in mind, we first examined the effect of fixed thresholds on clustering accuracy in a controlled dataset with known genome-sequence relationships. Subsequently, we introduce a data-driven approach aimed at more accurately clustering amplicon sequences to reflect their true source organisms.

### Clusters based solely on sequence similarity do not resemble original genomes in a synthetic community

In order to study the relationship between genomes and their corresponding *16S* gene sequences using fixed thresholds, we utilize amplicon sequencing data from an experiment that uses a synthetic community (SynCom) of 134 microbes, all of which were isolated from plant roots grown in natural soil and with sequenced genomes (Selten et al., 2024a). This dataset originates from a plant root colonisation experiment in which the SynCom members are exposed to the roots of different plant hosts and varying nutritional conditions. First, we assessed the number of different V3-V4 *16S* variable region sequences associated with these isolates. Among them, 65 isolates contain more than one V3-V4 sequence and were therefore named multiple-variant (MV) isolates. The remaining isolates are considered single-variant (SV) isolates. Overall, approximately 63% of genomes share at least one *16S* variant with any other. Many of these variants appear in few genomes and are shared by small and isolated groups of microbes, although a considerable number of them show a fuzzy distribution across genomes (**Figure 1B**), which translates into a large group of isolates sharing variants in a complex pattern (**Figure 1C, D**). To further study the impact of different clustering cut-offs on the identity of the resulting clusters, we calculated all pair-wise sequence similarities and clustered all sequences into OTUs from 100% to 90% similarity (**Figure 1E**). For this, we used the *k*-mer-based sequence similarity (*k* = 5) instead of classical alignments, as the former are faster to compute and represent local alignments accurately (**Supplementary Figure 1**). We named the resulting clusters *k*-mer-based OTUs, or *k*OTUs. As expected, the total number of *k*OTUs decreased with lower thresholds, and the number of chimeric *k*OTUs that merged multiple isolates, and even multiple species, increased (**Figure 1F, G**). At 100%, around 25% of all *k*OTUs already contains more than one single isolate and 20% of them contain sequences from more than one species (**Figure 1F**). The precise number of isolates and species merged into one single *k*OTU ranges from 1-10 for 100% similarity to 1-42 at 90% (**Figure 1G**). At the same time, around 60% of MV isolates are split into multiple *k*OTUs at 99%, and this is still 50% at 97% similarity (**Figure 1H**). In conclusion, and in agreement with previous studies, we did not find any “one-size-fits-all” threshold that could meaningfully cluster amplicons into units resembling the original genomes.

### Correlation in sequence abundances reflects latent intragenomic co-occurrence

Given the above-mentioned limitations, it seems highly advantageous to incorporate additional biologically-relevant information that guides the clustering process to accurately assemble genome-like clusters. In metagenomics, for example, commonly used algorithms leverage sequence co-abundances to facilitate the assignment of contigs to potential genomes (Alneberg et al., 2014; Kang et al., 2019) These strategies have the potential to enhance clustering decisions and align the results more closely with biological reality. For co-abundance to be useful at grouping variants into genome-like structures, correlations between similar variants should reflect their co-occurrence within genomes (intragenomic co-occurrence). That is, co-abundance should exhibit a measurable relationship with intragenomic co-occurrence (**Figure 2**). In the SynCom data, all variants show at least some degree of co-abundance with any other sequence, and groups of correlated variants emerge visibly (**Figure 2A**). Among these correlations, many might arise from biological interactions between the different underlying organisms and, consequently, confound the statistical association to physical co-occurrence within individual genomes. Therefore, we decided to discard all correlations between very distant sequences and to only consider co-abundances between sequences at least 90% similar (**Figure 2B**). Then, to gradually reduce the effect of confounding correlations, we applied increasingly higher similarity thresholds to examine the relationship between co-abundances and intragenomic co-occurrence patterns (**Figure 2C**) using Mantel tests with 9,999 permutations. The analysis revealed a significant association between co-abundance and co-occurrence among variants, and the strength of this association increased as the sequences considered were more similar (**Figure 2D**), potentially due to the increasing removal of confounding comparisons. Interestingly, when considering both positive and negative correlations (any non-zero correlation), the association of co-abundance with co-occurrence was around 1.4 to 1.8-fold higher than when only positive correlations are considered. This suggests that negative correlations from similar sequences also carry information about their co-occurrence in genomes. These negative correlations might represent cases of competitive exclusion between similar strains, in which shared variants co-occur with other strain-specific variants in the genomes of competing microbes, rendering the strain-specific ones negatively correlated. In a similar manner, we used a *χ^2^*-test to study whether correlation status of variants — e.g., variants correlated vs not correlated — was related to their co-occurrence status — e.g., variants co-occur vs do not co-occur — (**Figure 2E**). Consistently, we observed a significant enrichment of correlated variants when they also co-occur, and an emphasised statistical signal — i.e., *χ^2^*-statistic — as similarity between variants increased. Notably, both the Mantel and *χ^2^* analyses show a clear transition around the 95% similarity threshold. Below this value, many of the remaining sequence pairs still represent comparisons across different genomes, where correlations are largely shaped by ecological interactions between strains rather than by physical linkage. These ecology-driven correlations dilute the association with true intragenomic co-occurrence. Around 95%, most of these non-informative comparisons have been filtered out, and the remaining pairs are increasingly dominated by variants from the same genome or closely related strains. This shift leads to a sudden strengthening of the association between co-abundance and intragenomic co-occurrence, producing the upward inflection observed at this threshold.

**Figure 2.**
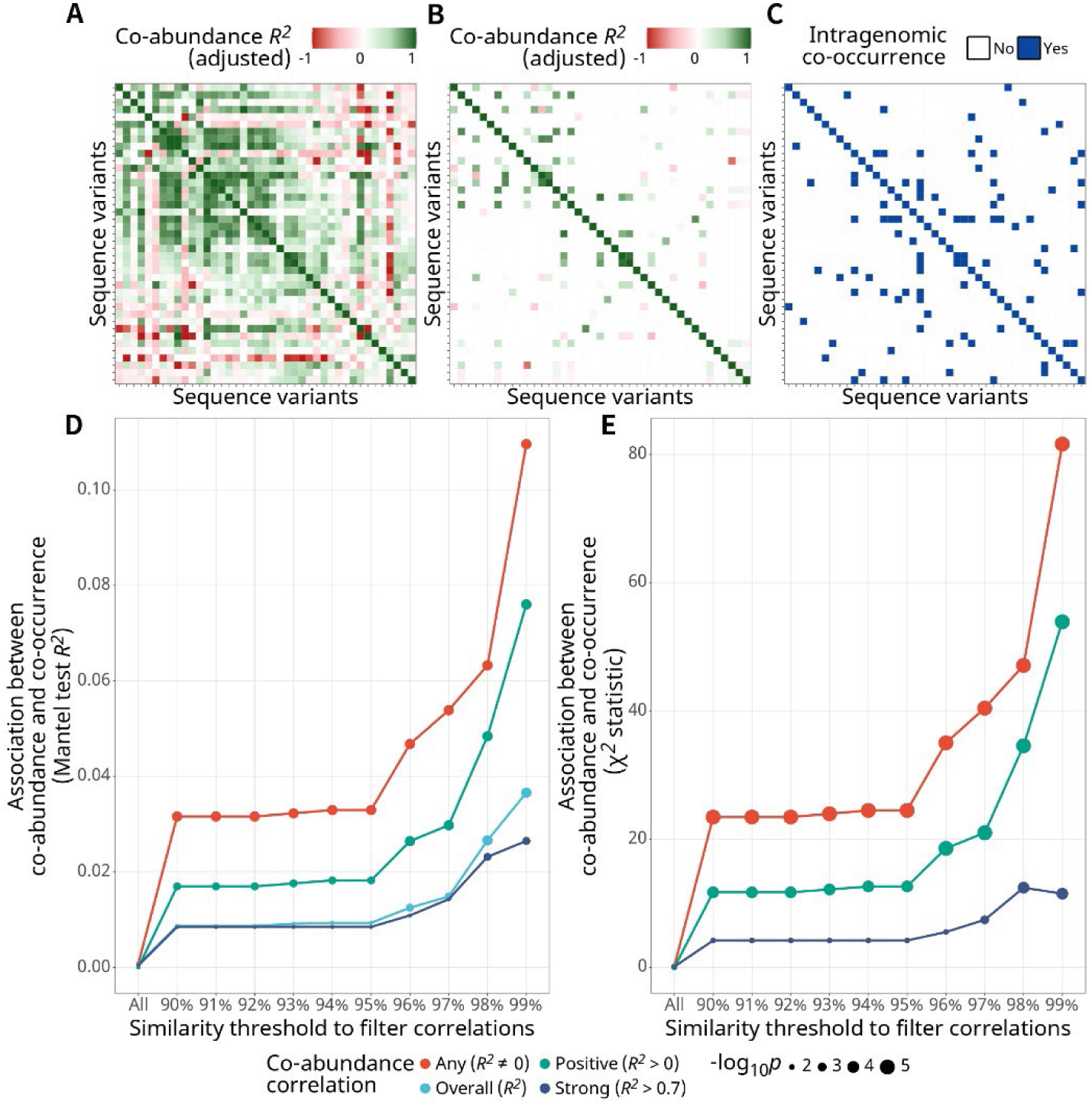
Using sequence similarity and co-abundance between variants can help enhance clustering. (A) Pearson co-abundance correlations (*R^2^*) of all unique sequences in the isolates. Rows and columns are ordered using Isomap seriation (Hahsler et al., 2025) applied to the co-abundance distance matrix. This procedure embeds the pairwise distances into a low-dimensional manifold and finds a 1-dimensional ordering that places similar variants adjacent to each other. (B) Same co-abundance matrix as in A, but only showing correlations for pairs of sequences at least 90% similar according to their *k*-mer similarity. Note how most of correlations in A came from distant variants, yet the matrix still depicts groups of somewhat similar sequences with strong correlations in the data, which could be used to cluster sequences into similarly-behaving groups of variants. (C) Intragenomic co-occurrence of all sequences in at least one genome, in the same seriation order as in A and B. (D) Line plot showing the result of a Mantel test comparing pair-wise abundance correlations in the SynCom data with their intragenomic co-occurrence pattern. The graph depicts the *R^2^* of the tests performed at increasingly higher thresholds. As the similarity between sequences increases, the test shows a higher association with their co-occurrence. (E) Line plot following the same strategy as in D in a χ^2^ test. Similar to the Mantel approach, the χ^2^ tests also reveal an increasing association between correlation status (correlated vs non-correlated) and the co-occurrence of variants within a genome as more similar sequences are considered. Any: all non-zero correlations are considered; overall: all correlation values are considered, unfiltered matrix; positive: positive correlations only; strong: only positive correlations above 0.7.

In conclusion, correlations of sequence abundance across samples indeed carry useful information about the occurrence of those sequences within the genomes in the community, and therefore, abundances can be leveraged to guide the clustering of *16S* gene sequences into biologically meaningful clusters.

### ASVNet: empirical OTUs from sequence similarity and co-abundance networks

We applied the principle described above into a novel algorithmic approach, which we named ASVNet — after <u>a</u>mplicon <u>s</u>equence <u>v</u>ariants <u>n</u>etworks —, aimed at reconstructing the correspondence between amplicon sequences and the organisms in a microbial community. ASVNet operates on the premises that different *16S* variants from the same genome will (1) be somewhat similar, and (2) strongly correlate across all samples. ASVNet follows these principles to integrate both sequence similarity and co-abundance information to infer empirical operational taxonomic units (eOTUs; **Supplementary Figure 2**). The concept of eOTUs and calling clusters based on both taxonomy and co-abundances was already followed by Probst and co-authors in the development of the PhyloChip technology (Probst et al., 2014). After hybridization of environmental DNA with the probes in a PhyloChip, eOTUs were inferred by finding correlated groups of taxonomically-related probes according to the fluorescence score across samples. Both for ASVNet and for PhyloChip eOTUs, the term ’empirical’ signifies that the resulting eOTUs are *ad hoc* and depend on the variation induced by or present in the experimental settings. Consequently, ASVNet requires sample-to-sample variation and variability induced by experimental conditions to increase the probability that any two individual microbes will exhibit a differential behaviour across samples, and therefore facilitate the clustering variants from an individual organism into a singular eOTU.

The algorithm starts by building a global sequence similarity network (SSN) of all variants, which is then partitioned into groups of similar sequences (SSN clusters). Subsequently, for each SSN cluster, ASVNet computes a sequence correlation network (SCN) based on co-abundances. Lastly, eOTU inference occurs in an iterative process that involves clustering each SCN into groups of sequences that are consistently correlated across samples. During the construction of the SCN, weak and noisy correlations are handled via either of two distinct modes. In the first and default mode, *empirical,* potentially spurious correlations are discarded by applying an *ad hoc* threshold. This threshold is determined by randomly sub-sampling pairwise correlations and selecting a specific percentile from the resulting distribution. The second mode, *regularised*, uses a regularised approach — elastic net regression — aimed at removing as many noisy and irrelevant correlations as possible while leaving the most meaningful ones, therefore enhancing resolution and inducing sparsity — the number of correlations that are turned into zeroes, and thus discarded. This mode requires setting a parameter, *α* — i.e., penalty mixture —, which controls the stringency of the method. When *α*=0, weak correlations are shrunk towards (but not made equal to) zero, whereas they are totally turned into zeroes and discarded when *α*=1.

### ASVNet eOTUs preserve isolate integrity better than kOTUs

To test ASVNet and search for the optimal configuration, we continued using the SynCom dataset as a ground truth, as the variability across nutritional conditions and plant hosts makes these data ideal to assess the quality of the algorithm. We searched for the best initial SSN threshold (*k*) and SCN construction mode, and we benchmarked the results from ASVNet against *k*OTUs at 97% and 99% similarity thresholds (**Figure 3**). We ran ASVNet with *k* = 90%, 92.5%, and 95%, in both *empirical* and *regularised* modes. For *regularised* runs, we tested multiple decreasing levels of stringency while still maintaining sparsity in the data, and therefore selected *α* values of 1, 0.9, 0.8, and 0.5. First, we explored the total number of clusters found by each method. As expected, *k*OTUs at 99% show the highest number of clusters (**Figure 3A**). Interestingly, ASVNet with an SSN threshold of *k* = 90% reported a total number of clusters that is very similar to the actual number of organisms in the SynCom, while applying higher SSN thresholds — i.e., *k* = 92.5% and *k* = 95% — increased the number of clusters approximately 2-fold with respect to the previous cut-off, but no dramatic differences are observed between different ASVNet modes. This shows that the selected SSN threshold strongly affects the total number of final clusters, with low values enabling abundance comparisons between very distant variants, and potentially resulting in more chimeric eOTUs.

**Figure 3.**
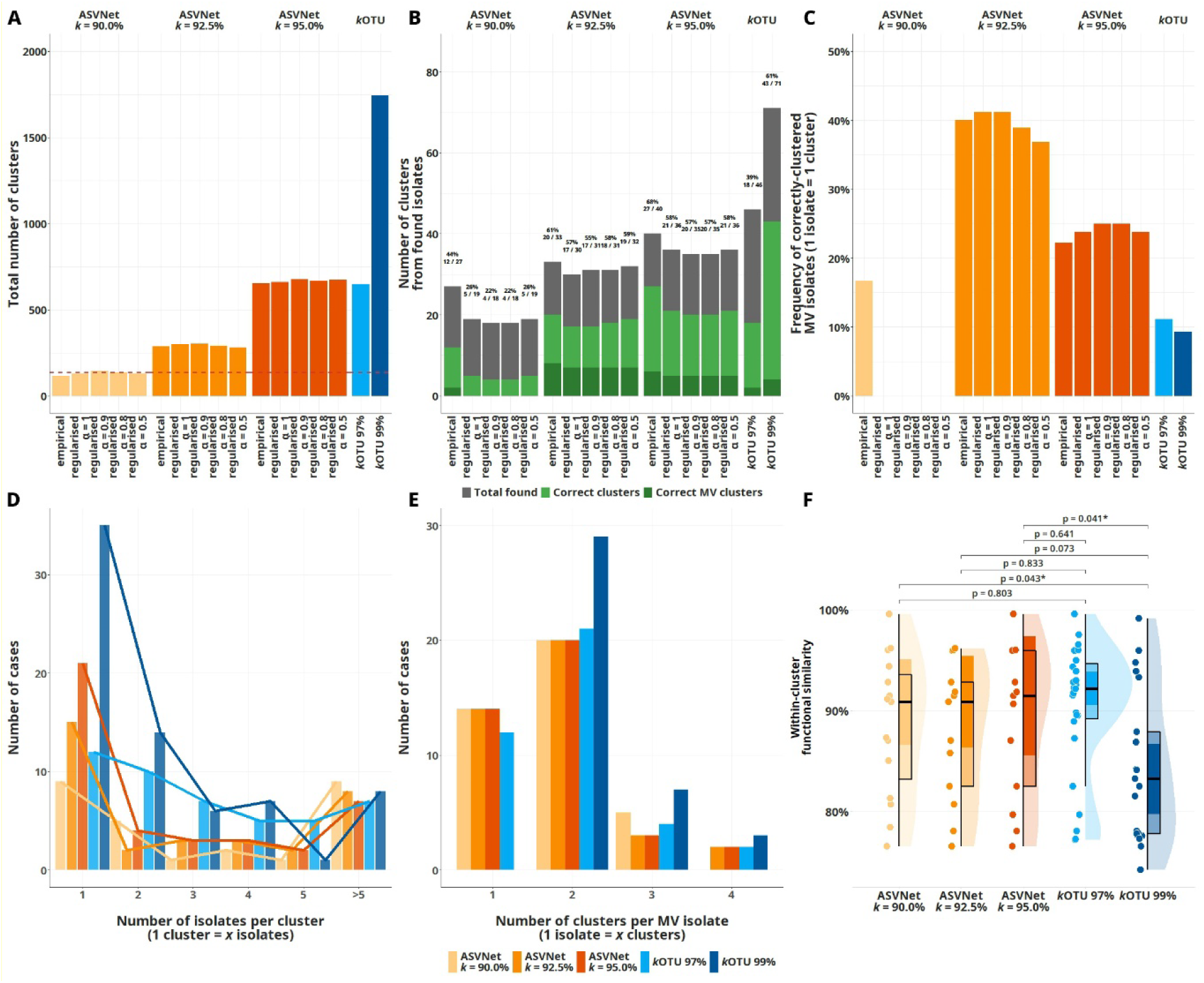
ASVNet resolves the SynCom dataset into more meaningful eOTUs. (A) Total number of clusters found by ASVNet and *k*OTUs at 97% and 99% sequence similarity. The dashed line marks the number of isolates in the experiment. (B) Number of clusters containing *rrn* sequences from the isolates in the microbial collection. The overlayed clear green bar depicts the total number of correctly-called isolates (both SV and MV isolates), and the dark green bar the total number of correctly-called multiple-variant (MV) isolates. Note that ASVNet finds the highest number of correct MV isolates. (C) Percentage of correctly-called MV isolates versus all correct clusters, showing that ASVNet after a cut-off of 92.5% has the highest ratio of correctly-called MV isolates. (D) Histogram of the number of isolates merged per cluster as a measure of the merging tendency of the algorithms. Note that ASVNet merges on average fewer isolates in every cluster. (E) Histogram of the number of clusters with sequences from MV isolates as a measure of the splitting tendency of the algorithms. ASVNet clusters MV isolates mostly as one or two eOTUs, while *k*OTU methods tend to split them more often into different *k*OTUs. (F) Median within-cluster functional similarity for incorrectly-called clusters, showing that clusters that do not correspond directly to an original *rrn* configuration are still biologically meaningful and have a higher within-cluster functional similarity than *k*OTUs at 99%. The darker shade on top of each boxplot shows the 95% confidence interval around the median value (notches).

Next, we compared the different methods for their ability to effectively cluster sequence variants from the SynCom isolates. We labelled clusters as correct if (1) it pointed to only one isolate and (2) all the observed sequences from that isolate were within that same cluster. Among all variant-isolate configurations, particularly resolving MV isolates poses a significant challenge, as the different variants should end up in the same eOTU. These cases are crucial for assessing whether the algorithm could effectively group sequences into groups resembling the original genomes. Notably, although *k*OTUs at 99% contained more correct clusters, those were mostly coming from SV isolates, meaning that, expectably, *k*OTUs at 99% splits better into individual sequences. However, the highest number of correctly-grouped MV isolates is achieved by ASVNet after SSN cutoffs *k* = 92.5% and 95% (**Figure 3B**), accounting for approximately 40% and 22% of all correct clusters, respectively (**Figure 3C**). In comparison, *k*OTUs had a much lower frequency of correct MV isolates, i.e., 9% for *k*OTUs at 97% and 10% for *k*OTUs at 99%. In general, the *regularised* mode did not perform better than the *empirical* counterpart, and thus we removed this mode for subsequent analyses.

Beyond perfect clusters, we also interrogated ASVNet and *k*OTUs for their tendency to generate chimeric clusters (**Figure 3D**; number of isolates within a cluster) and to split sequences from MV isolates into different clusters (**Figure 3E**; number of clusters per MV isolate). In general, chimeric ASVNet eOTUs agglomerate fewer isolates than *k*OTUs, and there are fewer cases of isolates split into multiple eOTUs than with *k*OTUs at 99%. Notably, chimeric *k*OTUs are based solely on similarity among sequences, while chimeric eOTUs may arise from isolates that behave similarly across samples, and therefore are a result of ecological similarity. To test whether chimeric eOTUs merge isolates in a more biologically meaningful manner, we evaluated the functional similarity among the organisms within chimeric OTUs. Specifically, we predicted the set of KEGG pathways for all organisms in each chimeric OTU and computed the Jaccard index for each pair-wise comparison. After this, we defined the median functional similarity of the chimeric cluster as the median of Jaccard index values for all comparisons. Both eOTUs and *k*OTUs at 97% show a median functional similarity within clusters of more than 90%, contrasting less than 85% in the case of chimeric *k*OTUs at 99% (**Figure 3F**). This suggest that chimeric eOTUs, although technically not pointing at single organisms, indeed merge isolates in a biologically more meaningful manner than *k*OTUs at 99% similarity.

### ASVNet eOTUs better resemble original isolates in complex datasets

We observed that ASVNet can successfully cluster sequence variants from MV SynCom isolates, and consequently displays enhanced performance compared to *k*OTUs across various SSN thresholds. However, despite the high complexity of this particular SynCom, this dataset is still relatively simple compared with real-world communities, and derives from a controlled environment with few organisms and a small number of different variants. Yet, the main objective of ASVNet is for it to be applied to natural community microbiome datasets. Such datasets typically exhibit high diversity and encompass numerous closely-related organisms. Organisms in which sequence variants co-occur in a more intricate pattern, and where many variants show significant sequence similarity to others. One of such complex environments is the plant rhizosphere, a dynamic environment formed by plant roots and the narrow region of surrounding soil under their direct influence (Poppeliers et al., 2023). We used two separate rhizosphere microbiome datasets to test ASVNet performance. The ‘*rhizosphere gradient experiment*’ data contains 96 samples consisting of microbial communities in different compartments of the natural rhizosphere, achieved through increasingly stronger washes aimed to enrich for the microbes on the plant roots (Poppeliers et al., 2024). The ‘*root time series experiment*’ data also contains 96 samples, all of them from the same rhizosphere compartment, but in this case the data represents a longitudinal study that follows the development of the microbial community in time (unpublished data). Both experiments were performed with natural soil extracted from the same location, from which a microbial collection had been previously established and all its genomes made available (Berendsen et al., 2018; Selten et al., 2024a, 2024b; Stringlis et al., 2018). Furthermore, this collection captured approximately 53% of all original microbes found in the *Arabidopsis* rhizosphere (Selten et al., 2024a), which makes this microbial collection a valuable benchmarking tool to assess the comparison of eOTUs with *k*OTUs. Following the same principles as with the SynCom dataset, we called eOTUs with ASVNet in *empirical* mode after applying the same three SSN thresholds and clustered all sequences into *k*OTUs at 97% and 99% for comparison (**Figure 4**). In contrast with the simpler SynCom data set, ASVNet reported many more eOTUs than *k*OTU methods in both rhizosphere sets for all SSN thresholds (**Figure 4A**). This suggests that ASVNet is affected by sample complexity, potentially due to a difficulty to resolve the clustering of the SCN with an increasing number of low-abundant microbes with few observations. Indeed, eOTUs tend to contain fewer sequences and with overall less abundance than *k*OTUs (**Supplementary Figure 3**), as previously observed with the SynCom data (**Figure 3D**), which may indicate that eOTUs more closely reflect the expected biological structure.

**Figure 4.**
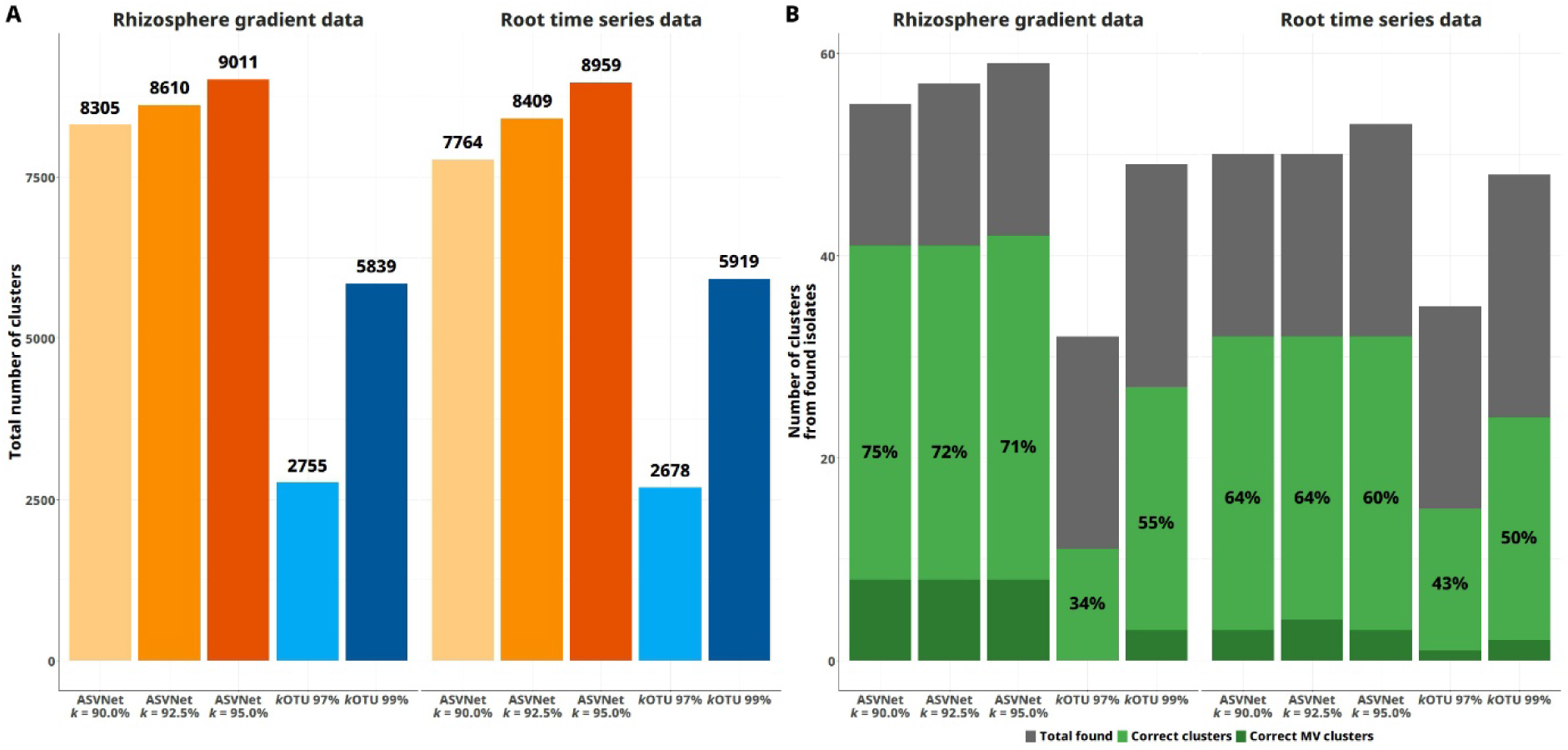
ASVNet finds more expected microbes in complex datasets. (A) Total number of clusters reported by ASVNet *empirical* and *k*OTUs at 97% and 99% similarity for both studies in this analysis. (B) Number of clusters containing *16S* gene sequences from the isolates in the microbial collection constructed from the same soil that was used in both studies. Note that the number of correct clusters, both for SV and MV isolates, is higher for ASVNet at any initial SSN threshold used.

For both datasets, we assessed the number of clusters containing *16S* gene sequence variants from the expected microbes. ASVNet eOTUs comprised the highest number of correctly classified isolates, approximately 70% of eOTUs, compared to 34% and 55% for *k*OTUs at 97% and 99%, respectively (**Figure 4B**). Importantly, eOTUs also contained the highest number of correctly classified MV microbes compared to *k*OTUs. These results show that integrating sequence similarity and co-abundance information in the clustering process helps construct eOTUs that better resemble the organisms in the ecosystem under study.

Altogether, our results show that co-abundance across samples can be used to solve the problem of clustering *16S* gene amplicon sequences into biologically-meaningful units. Moving forward, we aim to further refine ASVNet to address some of the important sources of complexity mentioned above, such as the assignment of sequences to more than one eOTU and the calculation of eOTU-adjusted abundances. So far, recent advances are achieving this when reference genomes are available (Selten et al., 2025), yet reconstructing genomes in reference-free settings remains a major challenge. In this sense, ASVNet could further benefit from incorporating advanced fuzzy clustering strategies and deep learning techniques to optimise the correspondence between variant co-abundance and the most-probable intragenomic co-occurrence structure. In addition, recent frameworks such as SyFi (Selten et al., 2025), together with the large body of publicly available microbial genomes and raw sequencing data, offer the possibility to build genome-informed, microbiome-independent eOTU references. These genome-derived eOTUs could serve as external priors to guide ASVNet in reconstructing organisms, particularly in settings where no sample-specific reference genomes are available. Ultimately, ASVNet lays the groundwork for future methodologies that aim to unveil the true composition of microbial communities through amplicon sequencing. By doing so, these future approaches have the potential to unlock deeper biological insights currently masked by multiple layers of complexity and embedded in both past and future amplicon sequence endeavours.

## Methods

### The ASVNet pipeline

#### Phase I: SSN clusters from original sequences

The input for ASVNet can be either raw FASTQ files from *16S* gene amplicon sequencing experiments or a sequence table with counts — e.g., as result of denoisers like DADA2 (Callahan et al., 2016). The first step in ASVNet is to cluster the sequences by sequence similarity. This avoids intensive calculations of all pair-wise correlations when sequences are not expected to come from the same genome. Pair-wise sequence comparisons are performed using the *k-*mer similarity (by default, *k* = 5), as it allows fast calculations. This similarity is based on the proportions of *k*-mers shared by each pair of sequences (Zielezinski et al., 2017) and aligns well with (local) alignment-based similarity (**Supplementary Figure 1**). After sequence comparisons, a low and permissive threshold is applied. The thresholds used in this study are based on the required threshold reported to collapse all sequences from *E. coli* into a single OTU, i.e., 94.75% (Schloss, 2021). Based on our results, the default value is set at 92.5%. After applying the threshold, the resulting SSN matrix is partitioned in clusters of broadly similar sequences with MCL v14.137 (Dongen, 2000), which uses Markov clustering for graph-based clustering. The MCL algorithm is used in various commonly used bioinformatics tools for protein homology clustering (to find orthologs) and to cluster genes from co-expression data. The resulting SSN clusters are subsequently input into the second phase of ASVNet.

#### Phase II: finding eOTUs

Due to technical imprecisions during amplification and sequencing, amplicon datasets accumulate many erroneous sequences that derive from the original true sequences. It is therefore expected that the initial correlations do not reflect biological reality in the samples. To account for this, eOTUs are inferred in each SSN cluster in a recursive process that starts by finding SCN clusters. The abundances for all sequences in a SCN cluster are merged into a single cluster and the process is repeated until no new strong correlations emerge or all sequences are merged (**Figure 5**). In a more detailed manner, the eOTU finding pipeline follows the next main steps:

**Figure 5.**
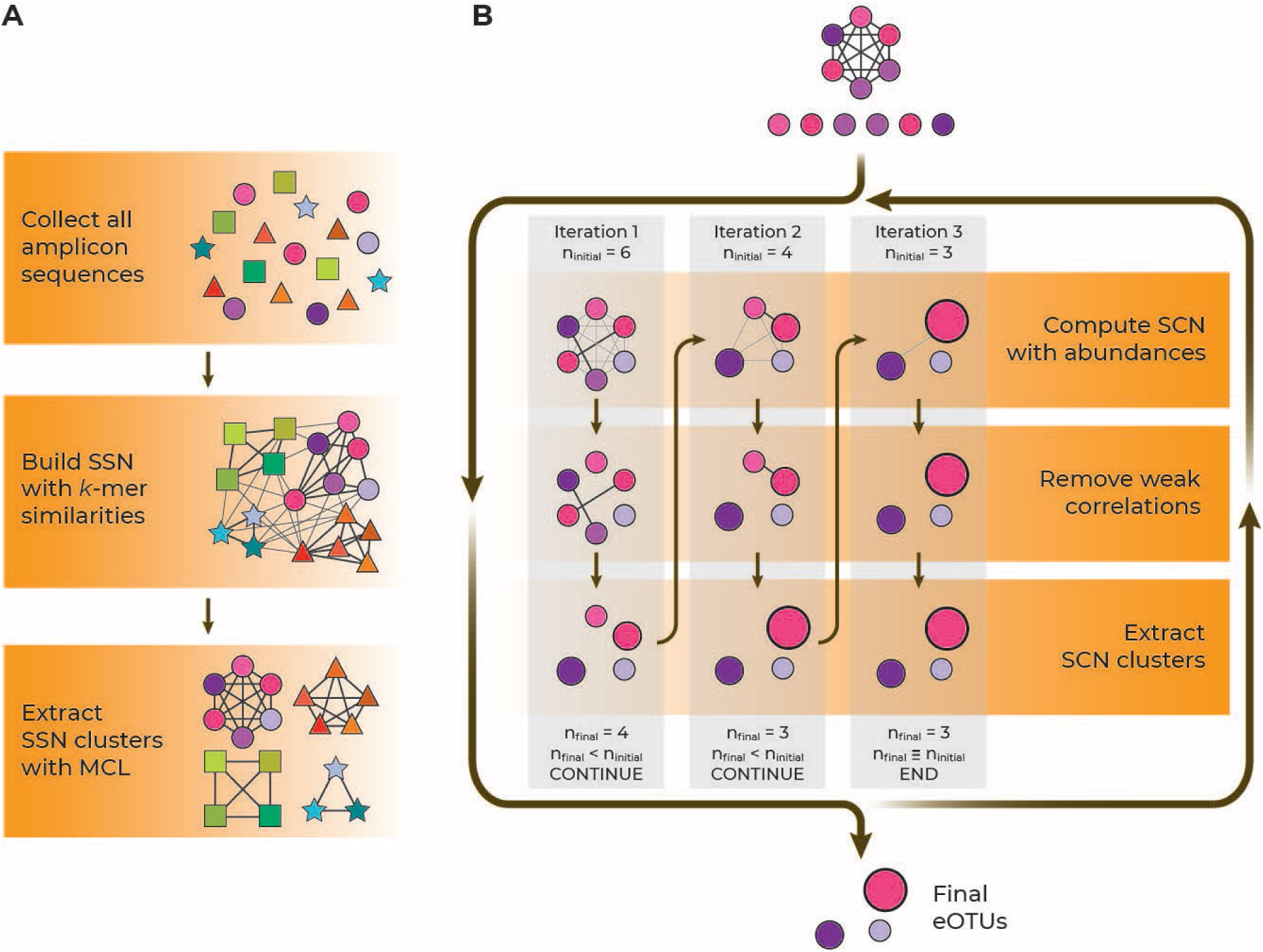
The ASVNet pipeline to infer eOTUs. (A) Phase I of the ASVNet pipeline. ASVNet collects amplicon sequences and builds a SSN using all pair-wise *k*-mer similarities, after which it partitions it into SCN clusters with the MCL algorithm. (B) Phase II of the pipeline. Per cluster of similar sequences, ASVNet builds the SCN using the correlations based on sequence abundances, filters low correlations, and finds clusters with MCL. ASVNet continues repeating this process with the clusters until no new correlations are found and eventually reports the resulting eOTUs.

**Step 1. Compute pair-wise linear correlations**. The initial counts are transformed with the centred log ratio (CLR) before calculating correlations as it has been previously suggested (Gloor et al., 2017). Both Pearson and Spearman correlations can be calculated, although Spearman is the default option.

**Step 2. Filter correlations**. Low correlations are removed using a rectifying function based on a logistic-adjusted rectified linear unit (ReLU). Briefly, the logistic-ReLU is constructed from bootstrapped correlations of the original matrix. These random correlations are used to calculate the correlation *p* at the *α^th^* percentile (by default, 99^th^ percentile). After this, a ReLU function with an inflection point at *p* is modified by a logistic-like curve that slowly moves into a slope of 1. This logistic-ReLU can then be used to turn all correlation values that can be observed by chance to zeroes, whilst it leaves the remaining correlation values unaffected.

The construction of the logistic-ReLU is done only once at the beginning of phase II but it is used throughout the pipeline in every iteration of the correlations to filter low values. The result is a sparser matrix that represents the SCN.

**Step 3. Find and evaluate clusters**. Clusters in the SCN are identified with MCL. In this step, ASVNet can be configured to find increasingly more fragmented clusters, and will select the most compact one with the lowest sum of internal negative correlations, assuming that eOTUs that better represent genomes will contain the fewest number of such internal negative correlations and best compactness.

**Step 4. Merge sequences in same cluster.** When the optimal clustering is found, the abundances inside each cluster are summed and a matrix with a reduced number of columns is returned. If there are still multiple clusters, the process is repeated from step 1 and will stop when no new correlations are found after filtering or all sequences are merged into a single cluster. When stopped, the final eOTUs are saved and ASVNet continues with the next SSN cluster.

In the *regularised* mode, ASVNet follows a slightly stricter path in which the linear correlations in step 1 are replaced by regularized regression (elastic net regression) with either an user-specified or a globally-optimized regularization penalty parameter λ, and a mixing parameter α (by default 1 for lasso regression). Elastic net regression continues iteratively until no more clusters are found, after which ASVNet continues the pipeline in the *empirical* mode using the previous clusters as initial clusters.

#### Processing reads and SSN construction

The amplicon sequencing samples from all three studies included here — i.e., ‘*SynCom experiment*’, ‘*rhizosphere gradient experiment*’, and ‘*root time series experiment*’ — were generated by amplifying the V3-V4 hypervariable region of the *rrn* gene (unpublished data). Briefly, raw reads were quality-filtered, adapter-trimmed, and merged using bbmerge from the BBMap v38.96 suite (Bushnell et al., 2017). Merged reads were dereplicated using the *derepFastq()* function from the DADA2 R package v1.21.0 (Callahan et al., 2016), which is transformed into a matrix of counts of unique reads. For each dataset, sequences with sufficient value variation (at least five different values) and prevalence (found in at least five samples), and samples with more than 1,000 counts, were kept for further processing. The SSN from the resulting sequences is constructed by calculating all pair-wise *k*-mer similarities with a parallelized version of the *kdistance()* function from the *kmer* R package (Wilkinson, 2018) and included in ASVNet as *kDistance()*. The result is the SSN matrix, which is further filtered with the different minimum similarity *k* thresholds, after which the SSN is partitioned into the SSN clusters with the MCL algorithm wrapper function *ssnMCL()* in ASVNet.

#### Code availability

ASVNet is available on a GitHub repository and can be accessed from https://github.com/jjsanchezgil/ASVNet. The code for figures in this manuscript is available at https://github.com/jjsanchezgil/ASVNet_figures_scripts.

## Acknowledgements

The authors want to thank Sanne Poppeliers, Gijs Selten, Bas E. Dutilh, Corné M.J. Pieterse, and rest of members of the Plant-Microbe Interactions lab for helpful discussions. This study was supported by the NWO Green II Grant no. ALWGR.2017.002 (J.J.S.G., R.d.J.) and the Novo Nordisk Foundation Grant no. NNF19SA0059362 (R.d.J.)

## Author contributions

**Juan J. Sánchez-Gil:** Conceptualization, Methodology, Software, Validation, Formal analysis, Investigation, Data curation, Visualization, Writing – Original Draft. **Ronnie de Jonge:** Conceptualization, Supervision, Project administration, Funding acquisition, Validation, Writing – Review and Editing.

## Supplementary Figures

**Supplementary Figure 1.**
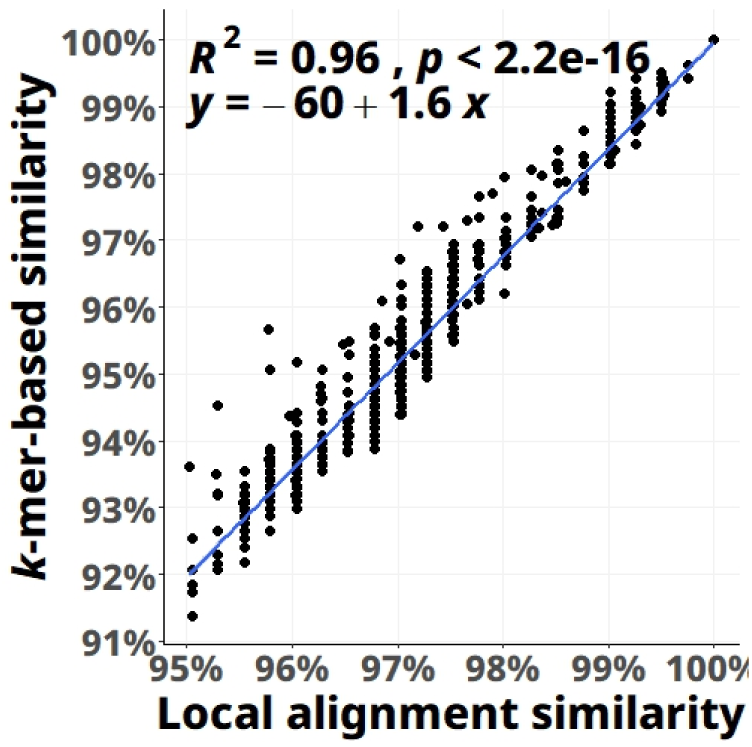
*K*-mer based similarity is very similar to local alignments. Scatter plot depicting the relationship between *k*-mer based (*k* = 5) and local alignment similarities of sequences in the SynCom dataset. The percent identity for local alignments was calculated as the proportion of identical positions divided by the length of the shorter sequence. The plot shows all pair-wise comparisons with an alignment similarity above 95%, extracted from a larger sample of 5,000 bootstrapped sequence pairs. Pearson’s *R^2^* for the correlation is shown in the graph, together with its *p*-value and the equation for the linear regression. The high *R^2^* and the low *p*-value denote a strong correlation between both similarity metrics.

**Supplementary Figure 2.**
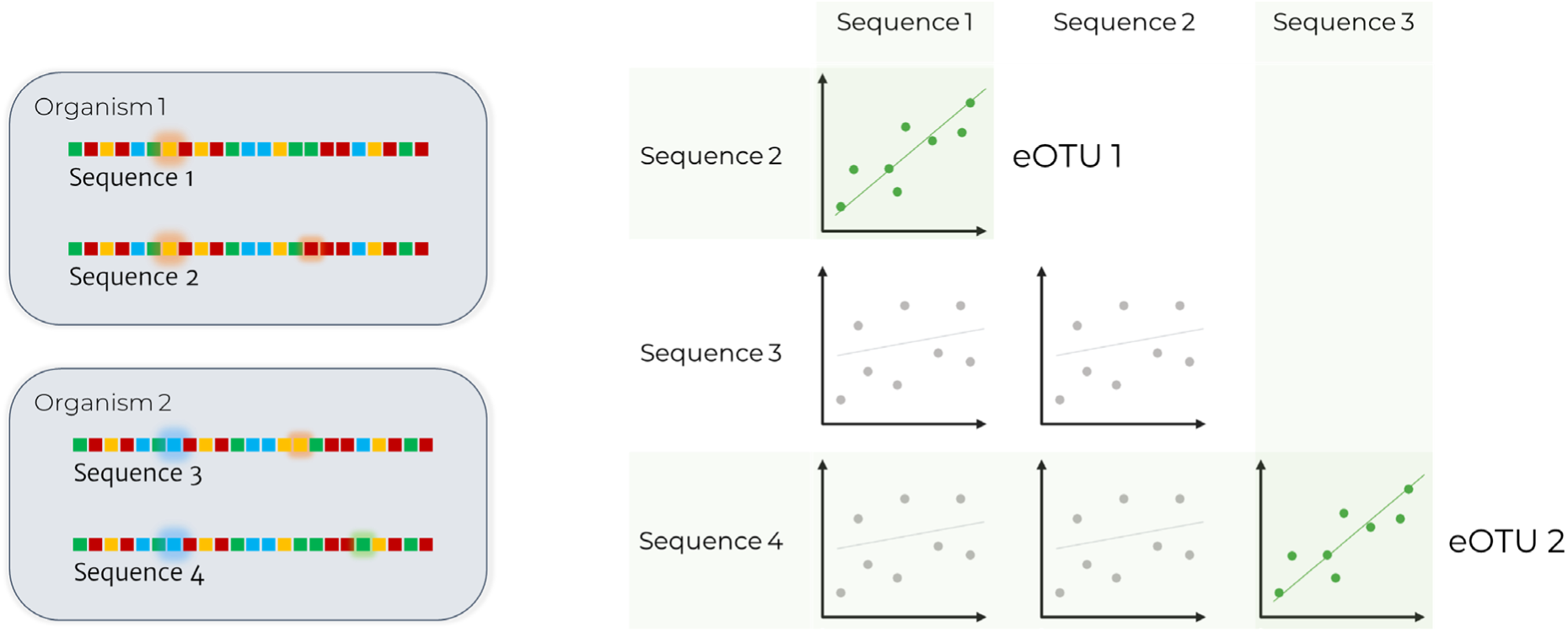
Conceptual overview of the rationale behind ASVNet. The left panel shows two example organisms, each containing two distinct *16S rRNA* gene copies (Sequences 1–4), illustrating intragenomic heterogeneity. Sequences from the same genome differ slightly but are more similar to each other than to sequences from other organisms. The right panel presents a toy correlogram of pairwise correlations in sequence abundances across samples. Variants from the same organism (Sequences 1 and 2 from Organism 1) show strong co-abundance correlations and are grouped into the same empirical OTU (eOTU 1). In contrast, sequences from different organisms (e.g., Sequence 3 and 4 from Organism 2) are less correlated and fall outside this eOTU. ASVNet integrates both sequence similarity and co-abundance patterns to group amplicon variants into eOTUs, aiming to reconstruct the underlying organismal composition of a microbial community.

**Supplementary Figure 3.**
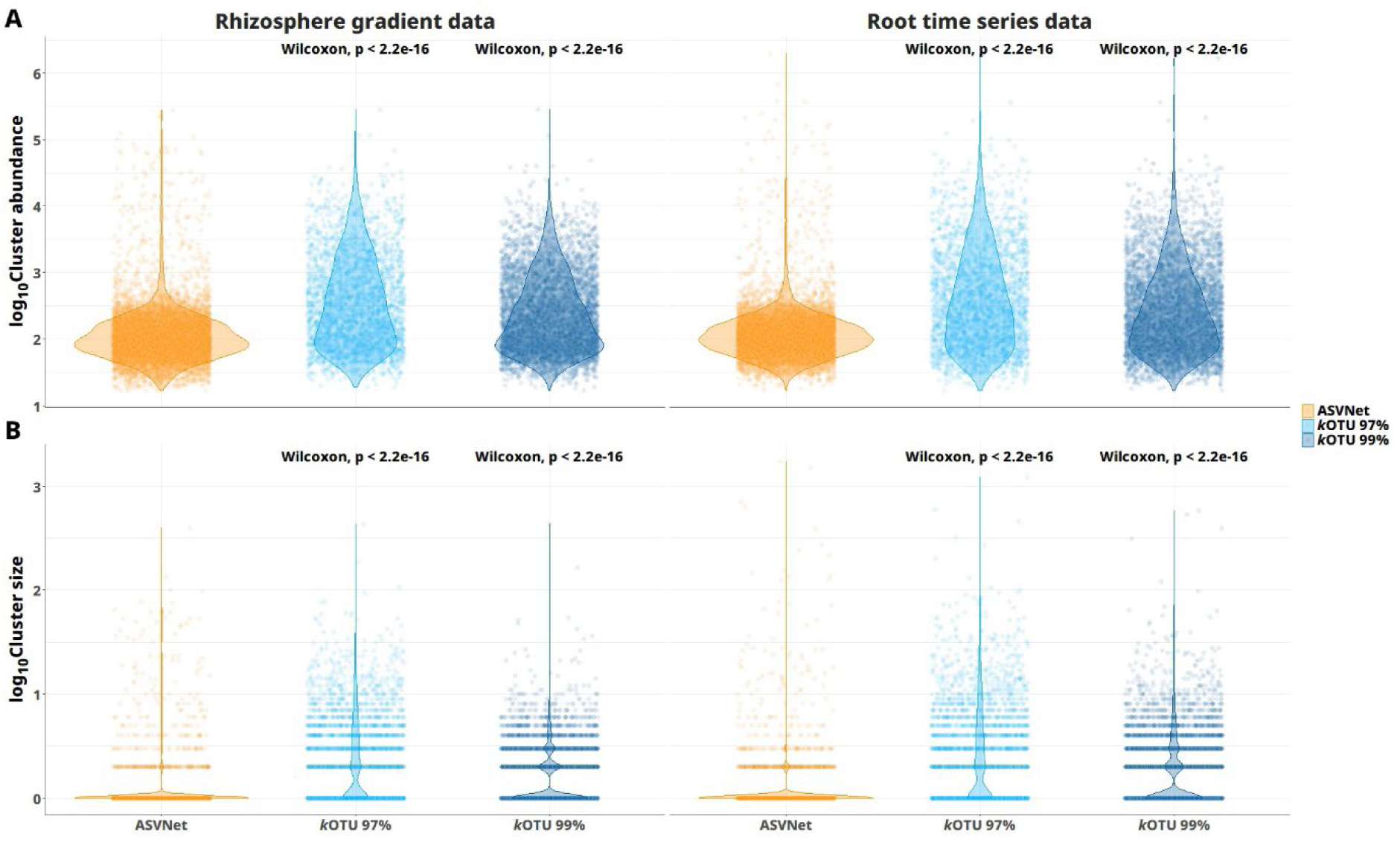
ASVNet eOTUs are less abundant and smaller clusters. Violin plots showing the log distribution of cluster abundances (A) and sizes (B) for ASVNet and *k*OTUs for the *‘rhizosphere gradient*’ data (left panels) and the *‘root time series data*’ (right panels). In every panel, the result of a Wilcoxon test comparing ASVNet to *k*OTU distributions is shown on top of each *k*OTU violin. For both datasets, ASVNet eOTUs tend to be less abundant and group fewer sequences than *k*OTUs at both similarity thresholds, as indicated by Wilcoxon tests.

