## Supplementary figures for "ASVNet: Inferring microbes from *16S rRNA* amplicon sequencing data"


#### Competing interests

This project received partial financial support from the private company Novozymes A/S, Denmark.

### Supplementary Figures

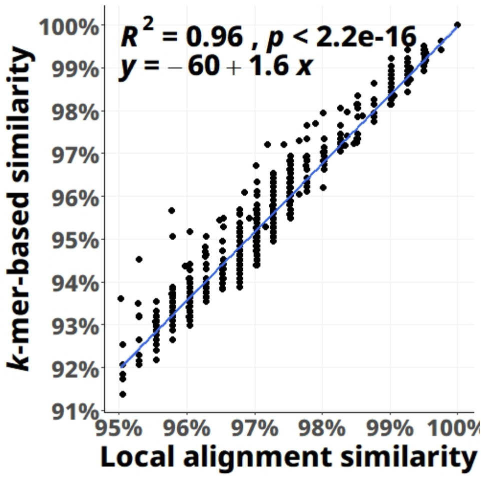


**Supplementary Figure 1. *K*-mer based similarity is very similar to local alignments.** Scatter plot depicting the relationship between *k*-mer based ($k$ = 5) and local alignment similarities of sequences in the SynCom dataset. The percent identity for local alignments was calculated as the proportion of identical positions divided by the length of the shorter sequence. The plot shows all pair-wise comparisons with an alignment similarity above 95%, extracted from a larger sample of 5,000 bootstrapped sequence pairs. Pearson's $R^2$ for the correlation is shown in the graph, together with its *p*-value and the equation for the linear regression. The high $R^2$ and the low *p*-value denote a strong correlation between both similarity metrics.

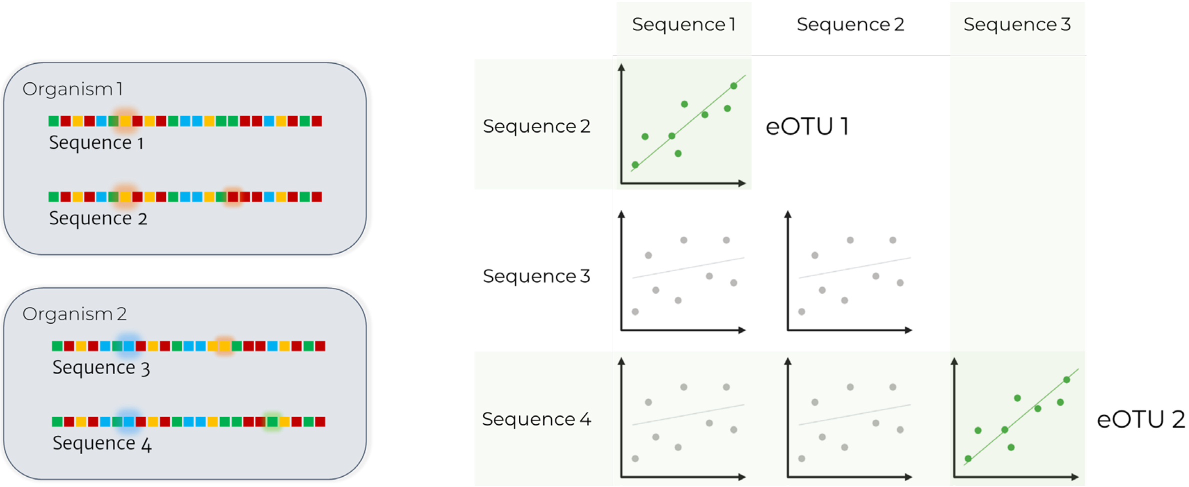


**Supplementary Figure 2. Conceptual overview of the rationale behind ASVNet.** The left panel shows two example organisms, each containing two distinct *16S rRNA* gene copies (Sequences 1–4), illustrating intragenomic heterogeneity. Sequences from the same genome differ slightly but are more similar to each other than to sequences from other organisms. The right panel presents a toy correlogram of pairwise correlations in sequence abundances across samples. Variants from the same organism (Sequences 1 and 2 from Organism 1) show strong co-abundance correlations and are grouped into the same empirical OTU (eOTU 1). In contrast, sequences from different organisms (e.g., Sequence 3 and 4 from Organism 2) are less correlated and fall outside this eOTU. ASVNet integrates both sequence similarity and co-abundance patterns to group amplicon variants into eOTUs, aiming to reconstruct the underlying organismal composition of a microbial community.

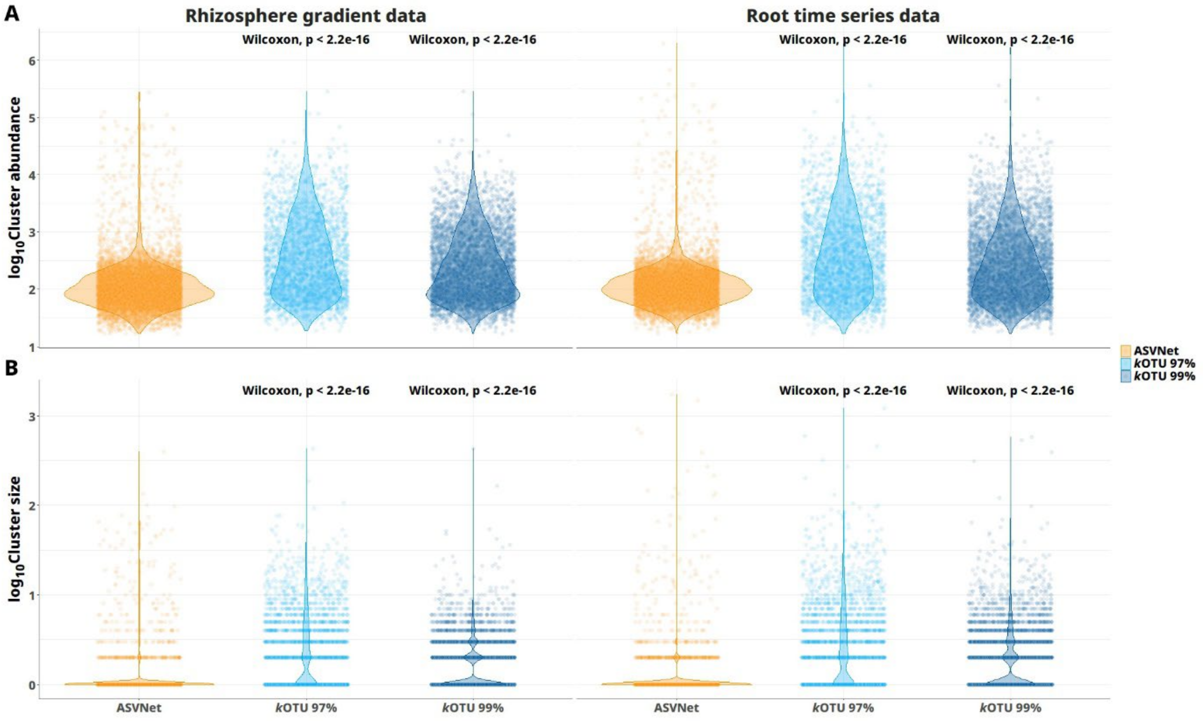


**Supplementary Figure 3. ASVNet eOTUs are less abundant and smaller clusters.** Violin plots showing the log distribution of cluster abundances (A) and sizes (B) for ASVNet and *k*OTUs for the '*rhizosphere gradient*' data (left panels) and the '*root time series data*' (right panels). In every panel, the result of a Wilcoxon test comparing ASVNet to *k*OTU distributions is shown on top of each *k*OTU violin. For both datasets, ASVNet eOTUs tend to be less abundant and group fewer sequences than *k*OTUs at both similarity thresholds, as indicated by Wilcoxon tests.
